# Using distant supervision to augment manually annotated data for relation extraction

**DOI:** 10.1101/626226

**Authors:** Peng Su, Gang Li, Cathy Wu, K. Vijay-Shanker

**Affiliations:** Department of Computer and Information Science, University of Delaware, Newark, Delaware, USA; Center for Bioinformatics and Computational Biology, University of Delaware, Newark, Delaware, USA

## Abstract

Significant progress has been made in applying deep learning on natural language processing tasks recently. However, deep learning models typically require a large amount of annotated training data while often only small labeled datasets are available for many natural language processing tasks in biomedical literature. Building large-size datasets for deep learning is expensive since it involves considerable human effort and usually requires domain expertise in specialized fields. In this work, we consider augmenting manually annotated data with large amounts of data using distant supervision. However, data obtained by distant supervision is often noisy, we first apply some heuristics to remove some of the incorrect annotations. Then using methods inspired from transfer learning, we show that the resulting models outperform models trained on the original manually annotated sets.

## Introduction

In recent years, deep learning has achieved notable results in several fields, and there is growing interest in applying deep learning for new tasks. With the explosive growth of text in the biomedical literature, applying natural language processing (NLP) techniques and deep learning to this field has attracted considerable attention. Relation extraction (RE) plays a key role in information extraction and aids the database curation for many disciplines [1] [2] [3]. The RE task is to identify relations between entities mentioned in natural language texts and its importance in biomedical domain stems in large part due to the fact that manual curation lags behind the growth in biomedical research literature. Developing high-performing systems to automatically extract relations from text is critical, and filling an important need.

There has been considerable effort invested in the extraction of different relations in BioNLP. It is fairly typical to cast relation extraction as a binary classification problem: where an instance comprising of a sentence and entities mentioned in the sentence are annotated as positive or negative depending on whether that sentence expresses a relation of interest among the marked entities. Many traditional (non-deep learning) machine learning methods have been applied on these problems (see e.g. [4] [5] [6] [7]) with most of them being feature-based or kernel-based methods. However, features/kernels have to be manually designed and their performance are not up to par with deep learning models when there is sufficient data.

Recently, deep learning methods show great advancement in various NLP tasks. Convolutional neural network and recurrent neural network are two well-studied types of deep learning architecture in NLP field. Promising results have been achieved by CNN model [8] [9] and current state-of-art CNN systems on relation extraction usually utilize refined architecture to incorporate more lexical and syntactic information. In [2], they applied piecewise max pooling process after convolutional layer to extract the structural features between the entities. The proposed method (piecewise CNN) exhibits superior performance compared with pure CNN. Peng et al. [10] proposed multiple channels in CNN to incorporate the syntactic dependency information and better capture longer distance dependencies. Also, RNN model shows its advantage on relation extraction, the model in [11] achieves state-of-the-art results on protein-protein interaction (PPI) task only using the word embedding as the input of LSTM model.

However, each new task requires its own annotated data for training the deep learning model. The annotation process of data needs considerable human effort to put a label on each data instance and often requires domain expertise, especially in specialized fields like Biomedicine. This issue is particularly onerous with deep learning since the models require setting of a large number of parameters and hence typically require large datasets. Currently, only small datasets are available for a number of tasks and this situation can hinder us from achieving the full potential of deep learning models. In order to alleviate the data limitation problem, Mintz et al. [12] first introduced the terminology of distant supervision (DS) and applied this technique to generate a large dataset for Freebase relation extraction, which assumes that a piece of text (often a sentence) expresses a relation between entities if these entities are related according to a known knowledge base. Before that, Craven et al. [13] already used the relation instances (tuples) gathered from some databases to label abstracts gathered from Medline, which pioneered the distant supervision method. Since then, distant supervision has been applied on many NLP tasks. Go et al. [14] applied distant supervision to automatically classify the sentiment of Twitter messages, and Surdanu et al. [15] used distant supervision approach for the TAC-KBP slot filling task. In the biomedical field, distant supervision has also been proven to be effective on extracting protein subcellular localizations [7] and microRNA-gene relations [16]. In the case of RE, distant supervision can be used to automatically obtain large training datasets using a knowledge base and large amounts of literature.

Noise in the labeling from distant supervision is a well-known problem and this labeling problem can adversely affect the performance of deep learning models [17]. To reduce the noise, many techniques have been proposed and the results show their effectiveness on the improving performance of DS-based models. One solution is to relax the originally strong assumption of DS, which assumes that all mentions of entity pair from the knowledge base express that relation. Riedel et al. [18] proposed the at-least-one assumption, which assumes at least one relation expression for entity pair from the DS holds, and built a multi-instance single-label model based on the DS data to reduce the noise. Then the work of [19] and [20] extended it to multi-instance multi-label model, which allows more than one label for each entity pair mention. At the same time, many other methods have also shown their advantages of reducing noise in DS data. Zheng et al. [7] introduced a threshold for the frequency of dependency paths among positive examples to filter out noisy examples. A novel generative model that directly models the heuristic labeling process of distant supervision was presented in [21]. Min et al. [22] proposed algorithm that learns from only positive and unlabeled data to alleviate the incomplete knowledge base problem. In the paper [23], the authors applied three heuristics (closest pairs, top trigger words, high-confidence patterns) to reduce the noise in the generated data, and demonstrated the improvement on performance. Like [23], we explore the behaviors of machine learning model on different DS-generated datasets with different noise reduction heuristics. Next, we will design methods to train models that use two data sets, the DS-obtained data and manually annotated (MA) data. Normally, DS data has been used by itself and not in combination with manually annotated data. In this work, we consider transfer learning for this purpose. Transfer learning is a technique where a model (often called the source model) developed for a task is reused as the starting point for training a second (target) model on another related task [24]. The hypothesis is that since the target model starts with learned knowledge from the source model, it will achieve better performance than the models trained from a random start. Transfer learning is proven to be effective to improve the performance (see, for example [25] and [26]). It has been applied on many tasks in natural language processing with good effect [27] [28].

In this work, we will test our methods using two well-known relation extraction tasks in biomedical field. The first is the protein-protein interaction extraction task [29], probably the most widely-studied relation extraction task in the BioNLP domain. Our experiments on this task use AIMed [30], a widely used benchmark corpus. To verify that our results generalize beyond this task, we consider second task; one of extracting protein subcellular localization task (PLOC) [31], which had previously been a focus of DS research in the BioNLP domain. The amount of human-labeled datasets available for this task is small and might not be sufficient for training deep models. We will evaluate our methods for this task on a recently available human-annotated corpus – LocText [32]. Also, for learning we consider previously proposed methods that have been successfully used for RE: we choose the piecewise CNN (PCNN) model [2] from CNN models and pick LSTM-based model proposed by [11] from RNN models.

We have conducted experiments to address different questions regarding the use of DS-data to augment manually annotated data. The first set of experiments consider developing DS data and training models on the raw DS data as well as after applying noise-reduction technique on the raw data. We also consider training models on the manually annotated data. Thus, we obtain three sets of models give us baselines whose performance provides context to compare and interpret the results of the next two sets of experiments.

The second set of experiments focus on the main concern of this work: using DS–derived data to augment manually annotated data to obtain larger amounts of training sets with the hopes of achieving better performance of deep learning models. Our next set of experiments considers alternate ways of using DS-derived and manually annotated data. We evaluate the effectiveness of trained models using a simple combination of the two data sets as well as by using two different ways of applying transfer learning.

Our motivation in this work is to supplement manually labeled data especially when it might not be sufficient for effectively training deep neural models. In our final set of experiments, we try to experimentally determine how much DS-derived data can compensate for small size of manually labeled data. Therefore we vary the size of the manually labeled data and see how the performance changes with the size of the manually annotated data set.

## Materials and Methods

### Experiments Conducted

In this section, we conduct several experiments to build and evaluate models based on DS data and manually annotated (MA) data. As mentioned earlier, we test our methods on two tasks, the protein-protein interaction task and the protein subcellular location task. The details of the development of the DS data for both tasks and the use of existing manually annotated benchmark sets for training and evaluation are discussed later in this section. We use two ways to combine these two types of data: pure data augmentation and transfer learning. In addition, we also explore the effect of using reduced amount of MA data after acquiring a large-sized DS-labeled dataset.

### Developing DS-trained Models

We start off by investigating how well the models trained on only automatically derived DS data. In [23], the logistic regression model performs better on noise-reduced DS data (DSNR). We conduct a similar exercise using the noise-reduction heuristics in conjunction with our deep learning models. Specifically, we will train the deep learning model on DS-obtained dataset, test the model on manually annotated data and then apply the same process on datasets obtained after applying different heuristics.

The human-labeled data for the two tasks can also be used to train the models. These models will also be evaluated on these labeled sets using 10-fold cross-validation. The models discussed here, i.e., the ones trained on DSO (original DS data), DSNR and MA can serve as our baseline models to evaluate the models discussed in the next subsection.

### Using DS and MA Data

After obtaining DS-labeled datasets, an obvious question is to consider how to combine it with the manually annotated dataset. The most straightforward way to combine two datasets is to simply take the union of DS data and MA data, and we will use it as a baseline here. We will also employ transfer learning as discussed below to combine DS-obtained and MA data in this paper.

Transfer learning focuses on storing knowledge gained while solving one problem and applying it to a different but related problem and has been proven effective for deep learning. Typically, a learned model for a task is used as a pre-trained source model and then used as a starting point for training on a dataset for a related task. In this paper, we will pre-train the model on (noise-reduced) DS datasets, then fine tune the model on MA training set to further adjust the parameters of the model. As we explained previously, the source model of transfer learning is a model from a similar task. Given the DS data may not exactly meet the guidelines used in developing the MA corpus, we can take the two as representing training data for closely related tasks.

In the pre-trained model, the learned knowledge of data stores in the hidden layers’ weights. These weights mean convolution filter (feature map) weights and the fully connected layers weights for CNN model, meanwhile mean recurrent cell weights and the fully connected layer weights in RNN model. Since the fully connected layer weights play the role of classifying the label of instance based acquired features in theory, convolution filter weights and recurrent cell weights contain the most important information learned from pre-training data. In this paper, we do not eliminate fully connected layers weights directly, since their functionality is not well studied. Instead, we design two options for transfer learning: 1). only transfer the convolution filter weights/recurrent cell weights; 2). transfer both convolution filter weights/recurrent cell weights and fully connected layer weights.

Fig 1 shows the pipeline of transfer learning model. When we use manually annotated data in both training and test process, we will perform cross validation to obtain the final results.

**Fig 1.**
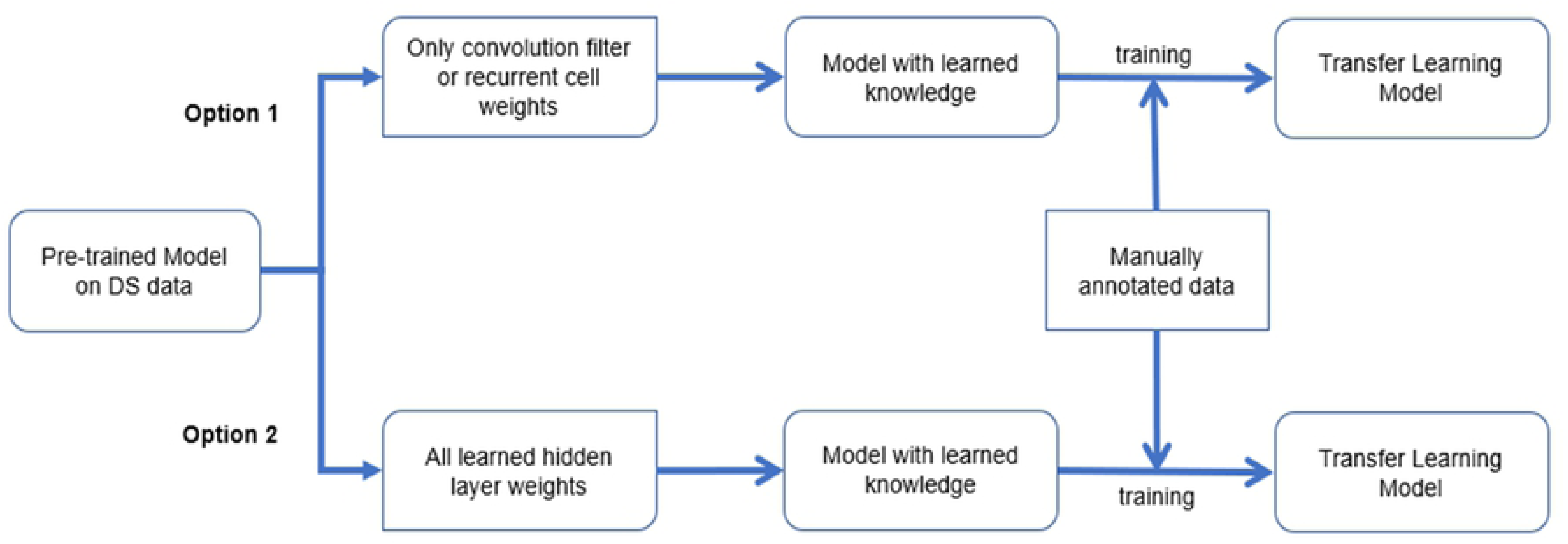
Pipeline of transfer learning model on both DS data and human-labeled data.

### Impact of Size of MA Training Data

The motivation for distant supervision is to have improved performance when there is only a limited amount of human-labeled data. Thus, it is worth examining the impact of the size of manually annotated data on the performance. For this set of experiments, we obtain transfer learning models pretrained on DS data and then use different sizes of manually annotated data to evaluate the dataset size effect. Specially, we will utilize 25%, 50% and 75% of the manually annotated data in the transfer learning training process to evaluate the performance of models.

## Experimental Setup Choices

### Neural Network Architectures

We have used both a CNN-based model as well as a RNN-based model in our experiments. The architectures were proposed in [2] and [11] respectively and shown to obtain excellent results.

In this section, we will briefly introduce the architecture of PCNN and LSTM model. Usually, the CNN model for classification problem contains: 1). convolution layer(s) to detect the local features; 2). pooling layer(s) to summarize the local features; 3). fully connected layer(s) to classify each category; 4) a softmax layer to output a normalized probability of each category. Fig 2 shows the structure of piecewise CNN model.

**Fig 2.**
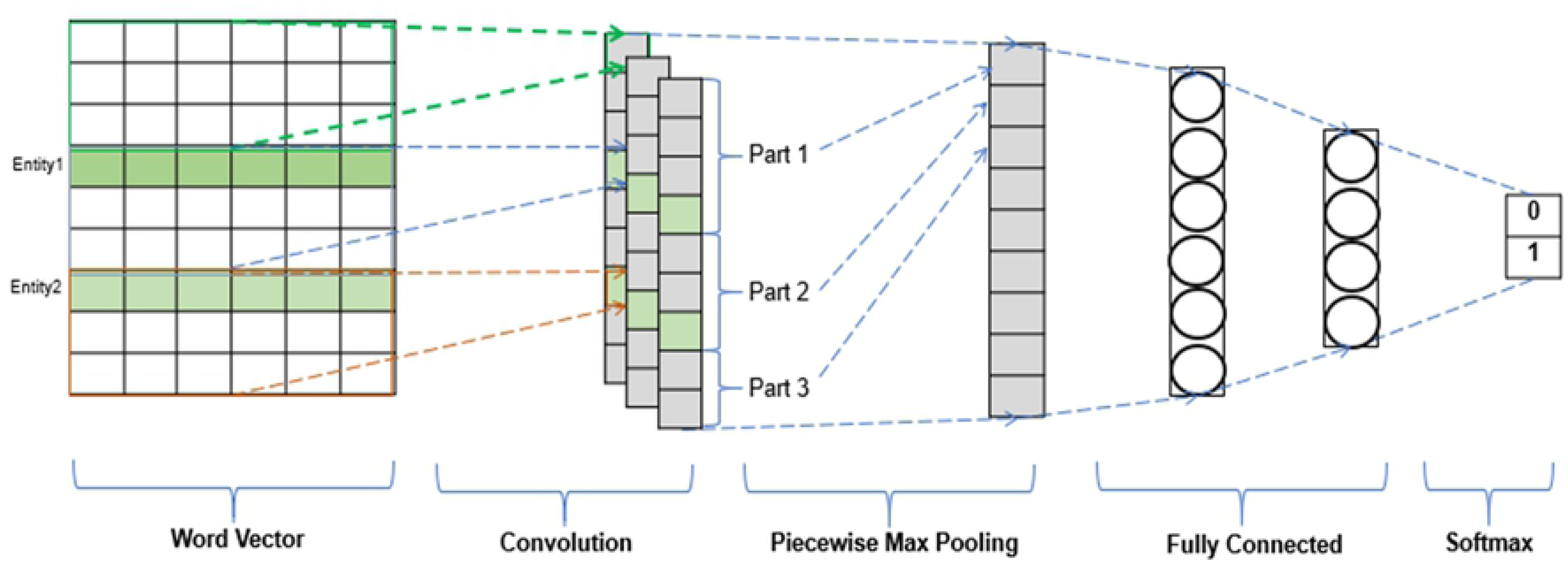
Structure of Piecewise CNN model.

The PCNN model is different with regular CNN models, whose max pooling is operated piecewisely based on the location of the entities in the sentence in order to include more structural information between the two entities. In this model, we divide the sentence into three parts using the entities as the segment points, and apply max pooling on these three parts separately. Let us take this sentence “We report an interaction between the human *PS*1_*PROTEIN*_ or PS2 hydrophilic loop and *Rab*11_*PROTEIN*_, a small GTPase belonging to the Ras-related superfamily” as an example, we will do maxing pooling on three parts: “We report an interaction between the human *PS*1_*PROTEIN*_ “, “or PS2 hydrophilic loop and *Rab*11_*PROTEIN*_ “, and “a small GTPase belonging to the Ras-related superfamily”. In this way, we will obtain three outputs for each sentence after max pooling and then we concatenate these three outputs as the output of max pooling.

Meanwhile the LSTM model has: 1). a embedding layer to generate the input sequence; 2). two recurrent layers (forward and backward) to model the sequence data in bidirectional way; 3). fully connected layer(s) to classify each category; 4). a softmax layer to output a normalized probability of each category. In Fig 3, we give the structure of LSTM model.

**Fig 3.**
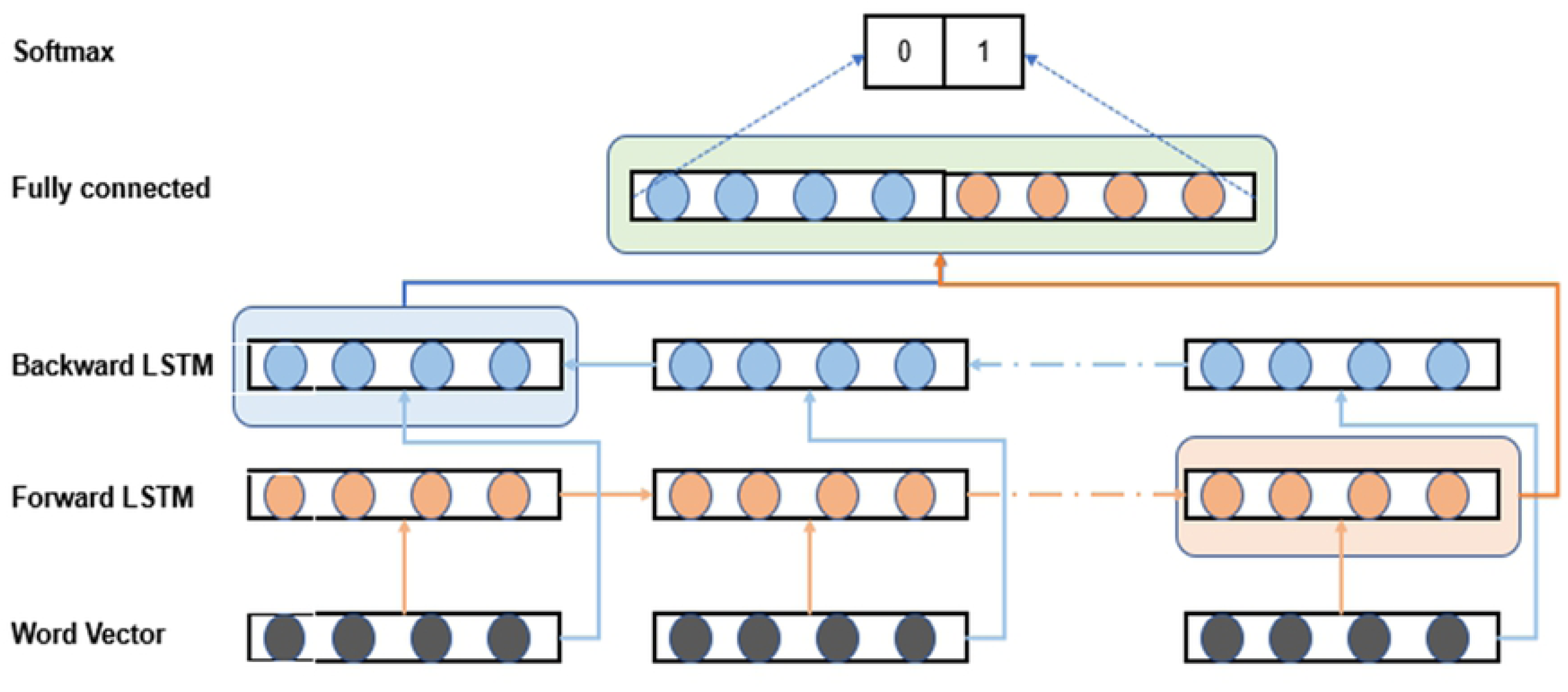
Structure of LSTM model.

### Input Representation

In this paper, we first represent each word by a word vector and then put all the word vectors in the same order with the sentence as our model input.

In this work, we found that both models perform better if we included more information about the input than what was included in the original papers. Specifically, we included in addition to the word embedding, POS tag, entity type, relative distance to two entities (see below), and incoming dependency relation in the word representation. The original PCNN model [2] only used word embedding and positional embedding (relative distance to two entities) to represent each word, while the LSTM model [11] only utilized word embedding of the sentence sequence as the input vector.

In this paper, we use the word embedding pre-trained on the PubMed using skip-gram model [33]. The dimension of each word embedding vector is 200. For POS tag and incoming dependency, we extract this information from the parse results of Bllip parser [34] and covert them to unique 10-dimension vectors. The relative distance to entities (to entity 1 (*d*_1_) and to entity 2 (*d*_2_)) is calculated by counting the words between the target word and the entities and the distance will be marked as negative if a word appears at the left side of the entity. After acquiring the distance numbers, we will map each number to unique 5-dimension vector. From the perspective of entity type, all the words in a sentence could be divided into four types: ENTITY1, ENTITY2, ENTITY, O. ENTITY1 and ENTITY2 are the two interacting entities, ENTITY is used for the other entities in the sentence, and O stands for other words. We use one-hot vector to represent to this feature.

### Parameter Choices

We implement the models with Tensorflow, the maximum length of sentence is set to 100, which mean the longer sentences are pruned and the shorter sentences are padded with zeros. The learning rate is 0.001 for PCNN model. Also, we apply decayed learning rate on PCNN with 0.95 decay rate and 1000 decay steps. For LSTM, we utilize constant learning rate of 0.001. We also apply dropout in these two models with drop rate of 0.5 on convolution/recurrent layer(s), and drop rate of 0.2 on dense layers. The training epoch is 30 for the DS and mixed data (DS+MA) for both models, which is the compromise of between the performance on mixed data and training time (more training epochs achieve slightly better results but need longer training time). The training epoch on MA data and transfer learning MA data is 200 for PCNN and 100 for LSTM (LSTM is trained with less epochs since it needs more time to train). Plus, the window size for PCNN is 3.

### Distant Supervision

We now discuss the knowledge database and biomedical text source to generate the distantly labeled data automatically.

For PPI task, we use IntAct database as the interacting protein pairs database, which is a freely available, open source database system for molecular interaction data [35]. We choose UniProt database [36] as our distantly supervised database for protein subcellular localization relation, which is a freely accessible resource of protein sequence and functional information.

Medline contains abstracts for biomedical literature from around the world and it is our first choice of text source, we use it for protein subcellular localization task by randomly sampling 30,000 abstracts that contains at least one pair of protein and subcellular location within one sentence. As it is shown in [23], it gives us a skewed dataset for the PPI task– positive/negative ratio is 1: 7.4. In order to acquire more balanced positive and negative instances for PPI, we just use the literature found in the IntAct database as our text source (Positive:Negative=1:1.5).

### Noise Reduction Heuristics

Distant supervision labeling process is noisy. It can generate false positive instances since it will always label the sentences with two entities stored in the known relation database as positive regardless of what the sentence says. For example, the sentence “The interaction between **bICP0** and IRF7 correlates with reduced trans-activation of the IFN-beta promoter by **IRF7**.” will be wrongly labeled as positive by DS even though there is no relation between these two highlighted entities. Also, distant supervision can introduce false negative instances due to the incompleteness of the relation database being used. For instance, DS will label the sentence “RFX5 specifically interacts with histone deacetylase 2 (HDAC2) and the mammalian transcriptional repressor (mSin3B), whereas **RFX1** preferably interacts with HDAC1 and **mSin3A**.” as negative by mistake, but it is obviously positive.

We considered a number of heuristics that have been proposed for DS noise reduction. We eventually decided to use the ones chosen in the work of [23], as it obtained good results. These heuristics are Closest Pairs (CP) and Trigger Words (TW) heuristics applied on the positive instances and High-confidence Patterns (HP) heuristic applied on negative instances.

We find that the definition of trigger word in the original paper is the ‘verb’ that expresses the relation between two entities, but the related two entities do not have verbal trigger word in many cases in PLOC task. So we only apply heuristic CP on positive instances for the PLOC task. Since we only apply two heuristics on PLOC DS dataset, we further filter out the noise by choosing the top 20 location names based on their frequency.

### Evaluation Sets

AIMed [30] is a widely used benchmark dataset for PPI task, we will use it as our evaluation set for PPI. LocText corpus [32] will be our evaluation set for PLOC task, which is a well annotated dataset with tagtog tool [37]. Please see the last row of Table 1 for the statistics of these two corpora.

**Table 1.** DS data and test data statistics. Baseline: original DS-labeled data without any heuristic; CP: Apply closest pair heuristic on DS data; HP: Apply high-confidence pattern heuristic on DS data; CP+TW: Apply closest pair and trigger word heuristics on DS data; CP+HP: Apply closest pair and high-confidence pattern heuristics on DS data; CP+TW+HP: Apply closest pair, trigger word and high-confidence pattern heuristics on DS data.

| PPI |  |  | PLOC |  |  |
| --- | --- | --- | --- | --- | --- |
| Dataset | Positive# | Negative# | Dataset | Positive# | Negative# |
| RAW | 54,170 | 82,517 | RAW | 19,654 | 32,254 |
| CP | 38,644 | 82,517 | CP | 15,519 | 32,254 |
| HP | 54,170 | 77,559 | HP | 19,654 | 30,206 |
| CP+TW | 25,294 | 82,517 | CP+TW | - | - |
| CP+HP | 38,644 | 77,559 | CP+HP | 15,519 | 30,206 |
| CP+TW+HP | 25,294 | 77,559 | CP+TW+HP | - | - |
| AIMed | 1,000 | 4,834 | LocText | 351 | 338 |

### Corpus Creation

In this section, we will introduce the creation of different DS datasets for each task. Given required knowledge base and text source for distant supervision, the last thing we have to consider is to label all the entity names (protein and subcellular location names) in the biomedical literature.

For protein names, we utilize the output of GNormPlus [38], which is an end-to-end system that detect gene/protein names. For the subcellular location names, we use location names from UniProt as a dictionary to match the mentions in the Medline text.

The first row in Table 1 shows the number of positive and negative instances we used for DS data for the two tasks. The next 5 rows show the size after applying different heuristics and their combinations.

## Results and Discussion

Throughout this section, we use precision, recall, F1 score as measurement to evaluate the performance of deep learning models.

### Models Trained on DSO, DSNR and MA Corpora

Although we differ from [23] in the choice of models which used Logistic Regression and Naive Bayes, we observe the same type of patterns when using the noise-reduction heuristics to filter the raw DS set (DSO). Other than some minor differences (e.g. precision of CP+TW+HP), the performance of the deep learning models are noticeably higher here.

The model built on noise-reduced DS data should achieve better performance as the heuristics on positive and negative will improve the precision and recall respectively. The noise in positive instances will make the model predict the negative ones as positive, so removing noise in positive instances will bring false positive rate down – precision improves. The noise in negative instances will lead the model to predict the positive ones as negative, so reducing noise in negative instances will make false negative rate decrease – recall increases.

Fig 4 shows the results of learning of the two types of neural network architectures on the two tasks. As is to be expected, precision improves with the use of CP, with the increase most noticeable in the PPI task case. Despite the drop in recall in LSTM-PPI combination, the F1 score improves in all four cases.

**Fig 4.**
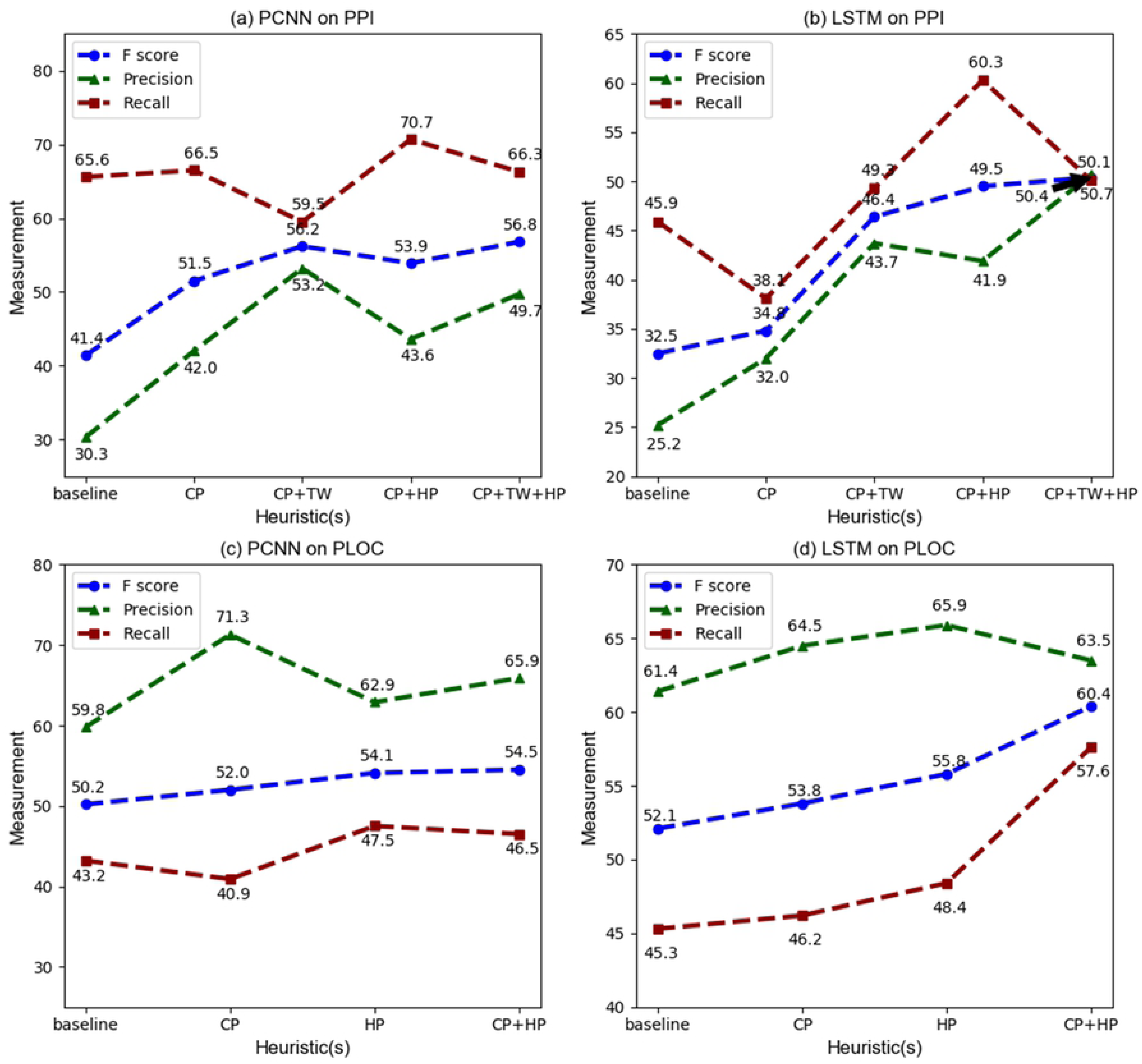
Performance of models built on DS data.

Next, the application of TW is considered and the expected increase in precision is noticed in both PPI graphs. As we discussed before, it is not proper to apply TW heuristic on PLOC dataset. In the PLOC case, we only see the improvement of precision on the use of CP heuristic.

As expected, the addition of HP boosts the recall in all four cases. Thus, we see that the addition of these noise-reduction heuristics helps boost the performance with F1 score showing an increase of 4% to 18%. In fact, in the case of PCNN on the PPI data, the performance on AIMed with no supervised learning is comparable to leading results obtained previously prior to the use of neural network models [10]. While the PCNN model obtains better results on the PPI task, the LSTM-based model performs better on the PLOC task, when trained on DS (with noise reduction) data.

Finally, we report the results for the same four combinations but this time using manually annotated data for training. As noted earlier, these results are based on 10-fold cross validation on the manually annotated sets for the two tasks. Row *Model*_*MA*_ of Table 2 and Table 3 shows the performance of the regular supervised learned models. Notice that the PCNN model achieves better F1 scores due to better recall results for both tasks, although the LSTM model has higher precision. Supervised learning on MA data improves the F1 score between 13% to 24% over noise-reduced DS trained model.

**Table 2.** Results of deel learning models on PPI. *Model*_*DSNR*_ : model built on noise-reduced DS data; *Model*_*MA*_: model built on manually annotated data (AIMed for PPI and LocText for PLOC); *Model*_*TLCON*_ : transfer learning using only the convolutional features; *Model*_*TLREC*_ : transfer learning using only the recurrent cell features; *Model*_*TLALL*_ : transfer learning using all the pretrained parameters. 10-fold cross validation is performed in these experiments.

| PCNN |  |  |  | LSTM |  |  |  |
| --- | --- | --- | --- | --- | --- | --- | --- |
| Model | Precision | Recall | F score | Model | Precision | Recall | F score |
| *Model<sub>DSNR</sub>* | 49.7 | 66.3 | 56.8 | *Model<sub>DSNR</sub>* | 50.7 | 50.1 | 50.4 |
| *Model<sub>MA</sub>* | 75.6 | 76.1 | 75.8 | *Model<sub>MA</sub>* | 78.1 | 69.7 | 73.7 |
| *Model<sub>Mix</sub>* | 65.0 | 68.2 | 66.5 | *Model<sub>Mix</sub>* | 64.2 | 54.9 | 59.2 |
| *Model<sub>TLCON</sub>* | 76.7 | 79.3 | 78.0 | *Model<sub>TLREC</sub>* | 78.3 | 74.5 | 76.4 |
| *Model<sub>TLALL</sub>* | 77.1 | 81.2 | **79.1** | *Model<sub>TLALL</sub>* | 78.9 | 74.7 | **76.8** |

**Table 3.** Results of deep learning models on PLOC.

| PCNN |  |  |  | LSTM |  |  |  |
| --- | --- | --- | --- | --- | --- | --- | --- |
| Model | Precision | Recall | F score | Model | Precision | Recall | F score |
| *Model<sub>DSNR</sub>* | 65.9 | 46.5 | 54.5 | *Model<sub>DSNR</sub>* | 63.5 | 57.6 | 60.4 |
| *Model<sub>MA</sub>* | 74.1 | 83.8 | 78.6 | *Model<sub>MA</sub>* | 76.0 | 70.5 | 73.2 |
| *Model<sub>Mix</sub>* | 71.4 | 69.9 | 70.6 | *Model<sub>Mix</sub>* | 72.7 | 63.7 | 67.9 |
| *Model<sub>TLCON</sub>* | 75.3 | 82.1 | 78.6 | *Model<sub>TLREC</sub>* | 79.2 | 78.3 | 78.8 |
| *Model<sub>TLALL</sub>* | 76.1 | 84.9 | **80.3** | *Model<sub>TLALL</sub>* | 80.3 | 78.4 | **79.4** |

### Combining DS and MA Data

This subsection is concerned with the core question of this work: how much improvement can we obtain by augmenting manually annotated data with (noise reduced) DS data. Table 2 and 3 show the results of various models. The first two rows in both tables are not based on the use of the combined data but instead repeat the results from previous subsection and provide the context for the new results using combined training data. The first row corresponds to the model *Model*_*DSNR*_ obtained by training on noise-reduced DS data, whereas the second row corresponds to the model *Model*_*MA*_ obtained by training on purely manually annotated data. As mentioned earlier, the pure data combination is the simplest way to utilize these two datasets, and hence we first consider the union of human-labeled data and (noise reduced) DS data. The third row of Table 2 and 3 shows the performance of the resulting trained models, designated *Model*_*Mix*_. The drop in the performance observed by comparing the second row suggests that simply taking the union of the instances on the two data sets may not be an appropriate way of augmenting the manually annotated data. Both precision and recall drop in all four cases. We hypothesize that the drop in performance might be due to some remaining noise in the DS data and/or that there might be some additional constraints in the manual annotation guidelines that might not be captured in the DS data.

Another way to combine the DS and human-labeled data is to use those pre-trained models as initial points, then further train the neural network models on manually annotated dataset, i.e. transfer learning. We have explored two options for transfer learning: 1). *TL*_*CON/REC*_: transfer learning with only convolution filter weights or recurrent cell weights; 2). *TL*_*ALL*_: transfer learning with all the weights of pre-trained model. The performance of these models trained in these manners is also shown in Table 2 and Table 3.

These two tables shows that both transfer learning models perform better than the models built on DS-labeled data as well as human-labeled dataset (AIMed and LocText). In fact, as hoped, the performance exceeds that of all other models, and obtains the best results ever. This implies that the deep learning models learn the knowledge in both DS and human-labeled data, and even though there may still be noise in DS data, the transfer learning process utilizes the human-labeled data to remedy the mistakes before and lead the learning in right direction in the second phase of model training. Thus, transfer learning is an effective way to make the best of DS labeled data and limited human-annotated data.

For the two options of transfer learning, we notice that the way of transferring all weights of pre-trained model obtains slightly better results overall. Thus, transferring all weights is our default way of transfer learning in our following experiments.

### Effect of Human-labeled Dataset Size

Any potential gains of the data augmentation method are more meaningful when the amount of available human-labeled dataset is not large. However, this is also a situation where any noise in DS derived data discrepancy between it and human-labeled data might hamper the effectiveness of data augmentation with DS data. This motivated the third set of experiments where we use noise-reduced DS data (and transfer learning) in conjunction with 25%, 50%, 75% of the human-labeled data in the model training process.

Fig 5 shows the F1 score corresponding to different sizes of human-annotated data, where 90% case corresponds to the results from previous subsection. The performance of the model obtained using transfer learning is shown and compared with those obtained with just the human-annotated (of the same size) data and with DS data. For example, training on 25% of AIMed data on the PPI task, the transfer learning method enables us to improve the performance by 10.6% and 22.2% using PCNN and LSTM respectively over the models of training on corresponding size of human-labeled data alone. We believe this shows reasonably good performance can be achieved with just 25% of manually labeled data using transfer learning, especially compared to using manually labeled data alone. Notice that with 25% of the data, the performance of the model trained on manually labeled data is worse than the model trained using DS data alone. The improvement using transfer learning narrows as the size of the human-labeled data increases. Improvement is also seen on the PLOC data, although the improvement is less than what was obtained for the PPI task. These results show that transfer learning and data augmentation approach always improves over the training on manual data alone, with the larger improvement shown when the size of human-labeled data is smaller, i.e., when there is limited human-labeled data, a situation which motivates this work.

**Fig 5.**
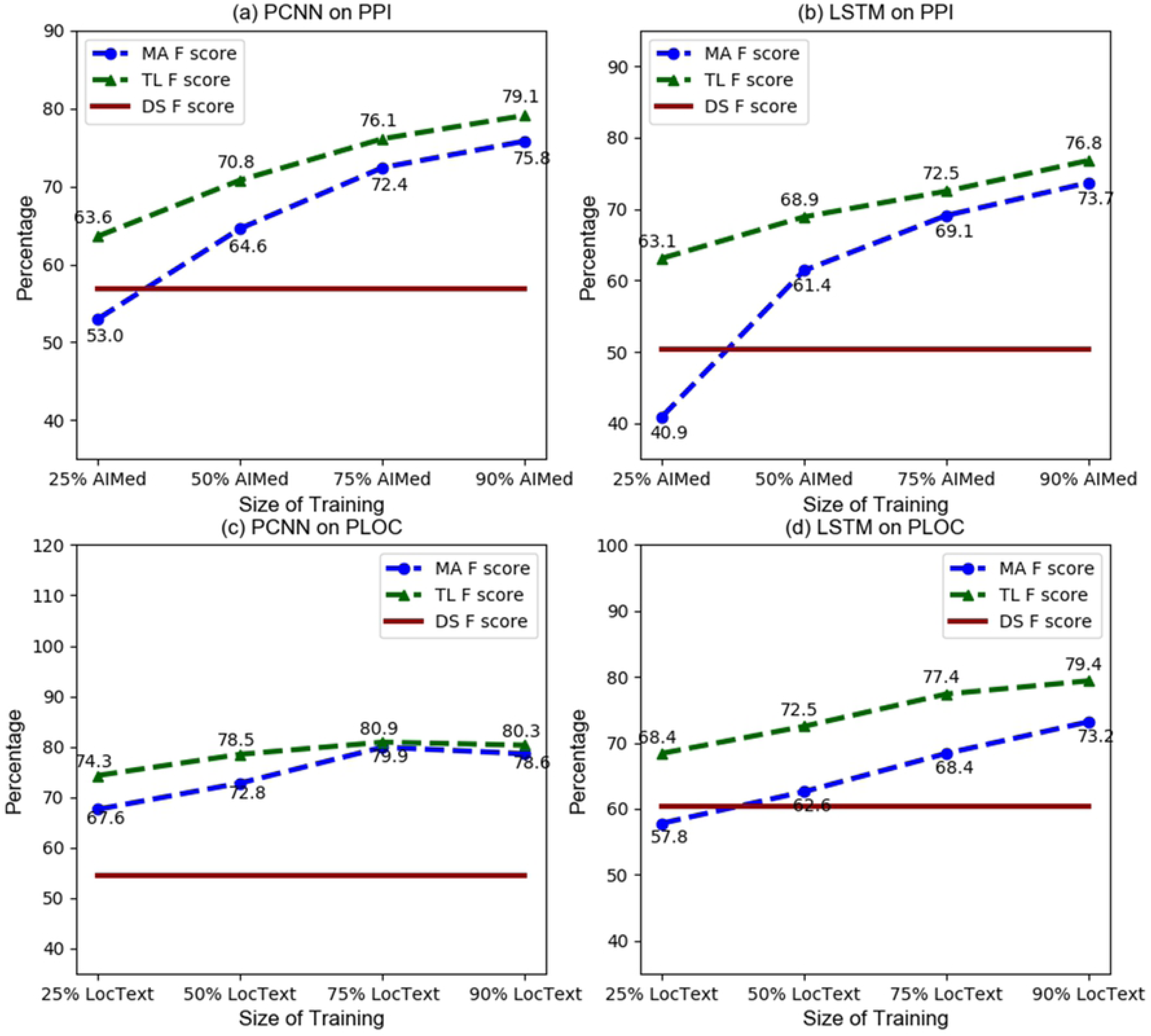
Trend of F score with different size MA data in transfer learning. MA F score means the F score acquired from models built on MA data only; TL F score means the F score acquired from models built on transfer learning; DS F score means the F score acquired from models built on DS data.

Fig 6 additionally presents the precision and recall numbers for more detailed analysis. For the PPI task with smaller amount of human-labeled data, most of the gains of transfer learning over just human-labeled data training are due to improvement in recall, although for LSTM-based model, the gains in precision are also substantially resulting in higher F1 score gain. With PLOC case, the gains in precision and recall are noticed.

**Fig 6.**
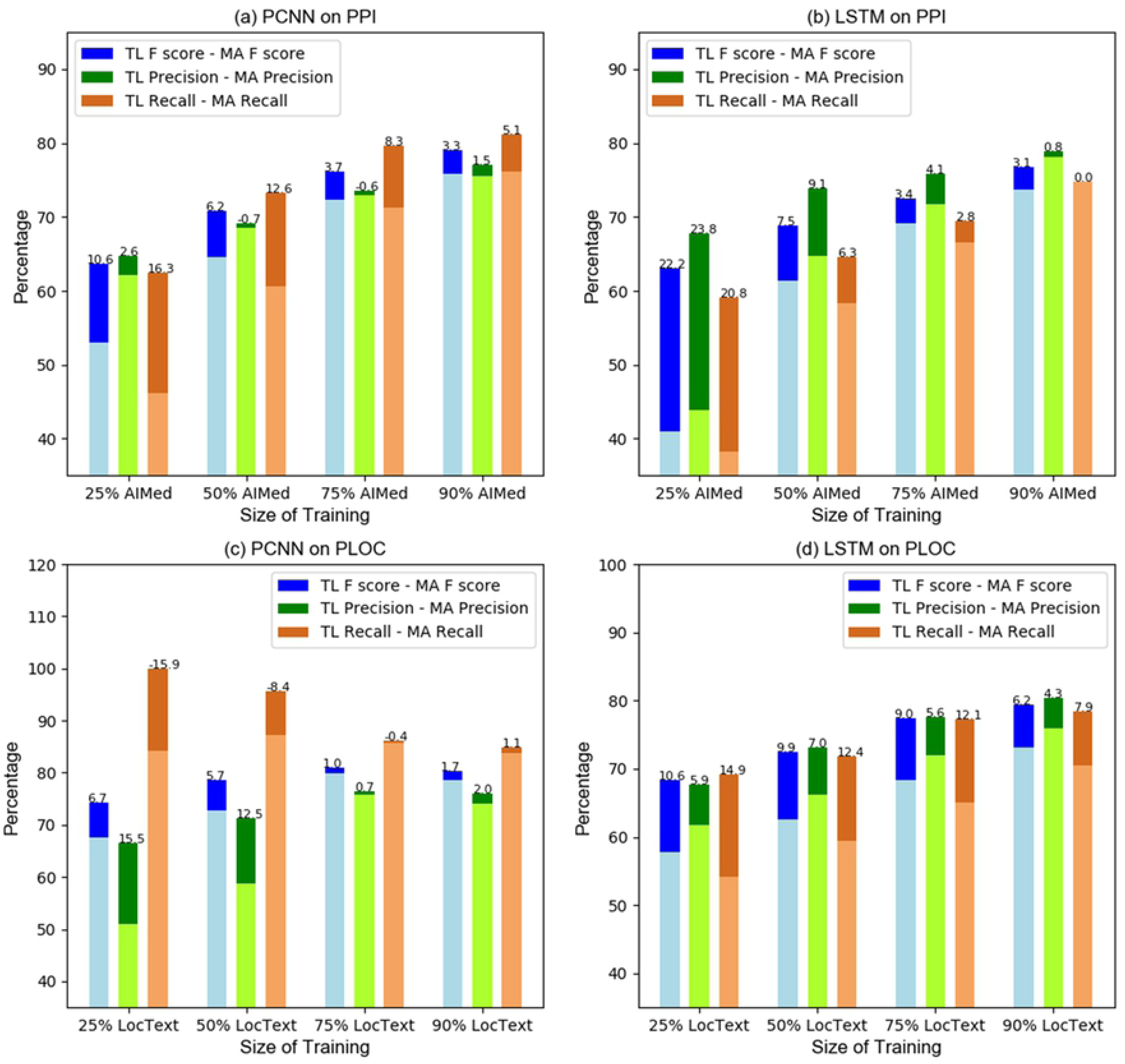
Size effect of human-labeled dataset. The number on each bar stands for the difference between None Transfer Learning and Transfer Learning model. Positive number means Transfer Learning improves the metric, while negative number means Transfer Learning deteriorates the metric.

## Conclusion

In order to improve the performance of deep learning models on small datasets, we have considered augmenting them with automatically obtained datasets using distant supervision. We show that some heuristics can be used to alleviate the well-known noisy annotation issue with distant supervision. Improvement of performance of both PCNN and LSTM models on both tasks is obtained.

Two methods of utilizing both DS data and manual data are discussed. Mixing DS data and human-labeled data to obtain the training data for deep learning model is the simplest way to combine data, but the performance does not show improvement over using human-labeled data alone. However, we show that the mechanism of transfer learning provides much better results than either of these two types of data individually.

We also explore the feasibility of reducing the size of manual data with the availability of large DS dataset. It can be seen that impact of transfer learning is much more beneficial when the manual data size is small (F score increased 10.6% when using 25% of AIMed). So when developing large human-labeled dataset is not feasible, applying transfer learning on DS data becomes more important.

These results are obtained for both types of deep learning models as well as both tasks, we plan to apply this technique on other relation extraction tasks. We will continue to pursue other heuristics to further reduce the noise in the automatic corpus creation with DS. Given the imbalance in the distribution of positive/negative instances in these datasets, we plan to conduct additional research to address this issue.

